# Differential effects of aneuploidy on growth and differentiation in human intestinal stem cells

**DOI:** 10.1101/2023.09.23.559117

**Authors:** Blake A. Johnson, Albert Z. Liu, Tianhao Bi, Yi Dong, Taibo Li, Dingjingyu Zhou, Akshay Narkar, Yufei Wu, Sean X. Sun, Tatianna C. Larman, Jin Zhu, Rong Li

## Abstract

Aneuploidy, a near ubiquitous genetic feature of tumors, is a context-dependent driver of cancer evolution; however, the mechanistic basis of this role remains unclear. Here, by inducing heterogeneous aneuploidy in non-transformed human colon organoids (colonoids), we investigate how the effects of aneuploidy on cell growth and differentiation may promote malignant transformation. Single-cell RNA sequencing reveals that the gene expression signature across over 100 unique aneuploid karyotypes is enriched with p53 responsive genes. The primary driver of p53 activation is karyotype complexity. Complex aneuploid cells with multiple unbalanced chromosomes activate p53 and undergo G1 cell-cycle arrest, independent of DNA damage and without evidence of senescence. By contrast, simple aneuploid cells with 1-3 chromosomes gained or lost continue to proliferate, demonstrated by single cell tracking in colonoids. Notably, simple aneuploid cells exhibit impaired differentiation when niche factors are withdrawn. These findings suggest that while complex aneuploid cells are eliminated from the normal epithelium due to p53 activation, simple aneuploid cells can escape this checkpoint and may contribute to niche factor-independent growth of cancer-initiating cells.

## Introduction

Colorectal cancers (CRC) are frequently aneuploid.^1^ Colon organoids, or colonoids, have been utilized to model tumor evolution.^2^ For example, cancer driver mutations have been introduced in colonoids to elucidate their individual contributions to niche factor-independent growth.^3,4^ Colonoids derived from colorectal cancers were used to demonstrate that continual chromosomal instability (CIN) generates novel aneuploid karyotypes and shapes genome evolution in colorectal cancer.^5^ Despite the growing evidence for aneuploidy’s role in cancer progression and the acquisition of drug resistance, it still remains elusive how aneuploidy, produced by erroneous mitosis, may be eliminated or tolerated in proliferating tissues and how it may contribute to the initiation of neoplasia. Whereas some studies suggested that the induction of aneuploidy activates p53 leading to growth arrest,^6–8^ others revealed that not all aneuploid cells activate p53.^9–11^ In our previous work, we demonstrated p53 activation is less robust in mouse colon and human mammary organoids when compared with certain cancer and immortalized cell lines,^12^ pointing to the possible importance of tissue context in the aneuploidy response and highlighting the need to investigate this question in cell models more closely mimicking normal human tissues.

In this study, we use colonoids derived from non-transformed human colon mucosa to assess the effects of aneuploidy on the fate of intestinal stem cells and the possible contribution of aneuploidy towards early tumorigenesis. Our data confirm that p53 activation is indeed a common consequence of aneuploidy, but it is primarily influenced by the extent of aneuploidy and largely independent of DNA damage. We demonstrate that some simple aneuploid cells continue to proliferate even in the absence of niche factors and may thus contribute to early cancer evolution.

## Results

### Induction of simple and complex aneuploidy in non-transformed human colonoids

We generated non-transformed colonoids from normal mucosa at the unaffected margin of a CRC surgical resection specimen. The mucosa used to generate these colonoids was histologically normal (Figure S1A). Colonoids grew at the expected rate and with similar morphology to previous reports.^13^ To induce diverse aneuploid cells with varied karyotype complexity, we treated the colonoids with different concentrations of reversine, an MPS1/TTK inhibitor^14^ (Figure S1B). Live imaging confirmed that reversine treatment accelerated mitosis and caused lagging chromosomes, chromatin bridges, and monopolar spindles (Figure S1C and S1D). Treatment with 0.25 µM reversine induced aneuploidy in 50% of cells with the vast majority of aneuploid cells having 1-3 aneuploid chromosomes (deviating from the 2n or 4n number), whereas 0.5 µM reversine induced aneuploidy in nearly 75% of cells generating a wider range of karyotypes including ∼40% cells having ≥ 4 unbalanced chromosomes (Figure 1A). We define simple aneuploidy as gain or loss of 1-3 chromosomes from the diploid or tetraploid number and complex aneuploidy as ≥ 4 gains or losses. As we did not expect MPS1 inhibition to generate triploidy, cells with chromosome numbers near 3n were categorized as complex aneuploidy. We define karyotype complexity as the mean difference from the diploid or tetraploid number normalized for ploidy for each condition and karyotype heterogeneity as the standard deviation of these differences.

**Figure 1.**
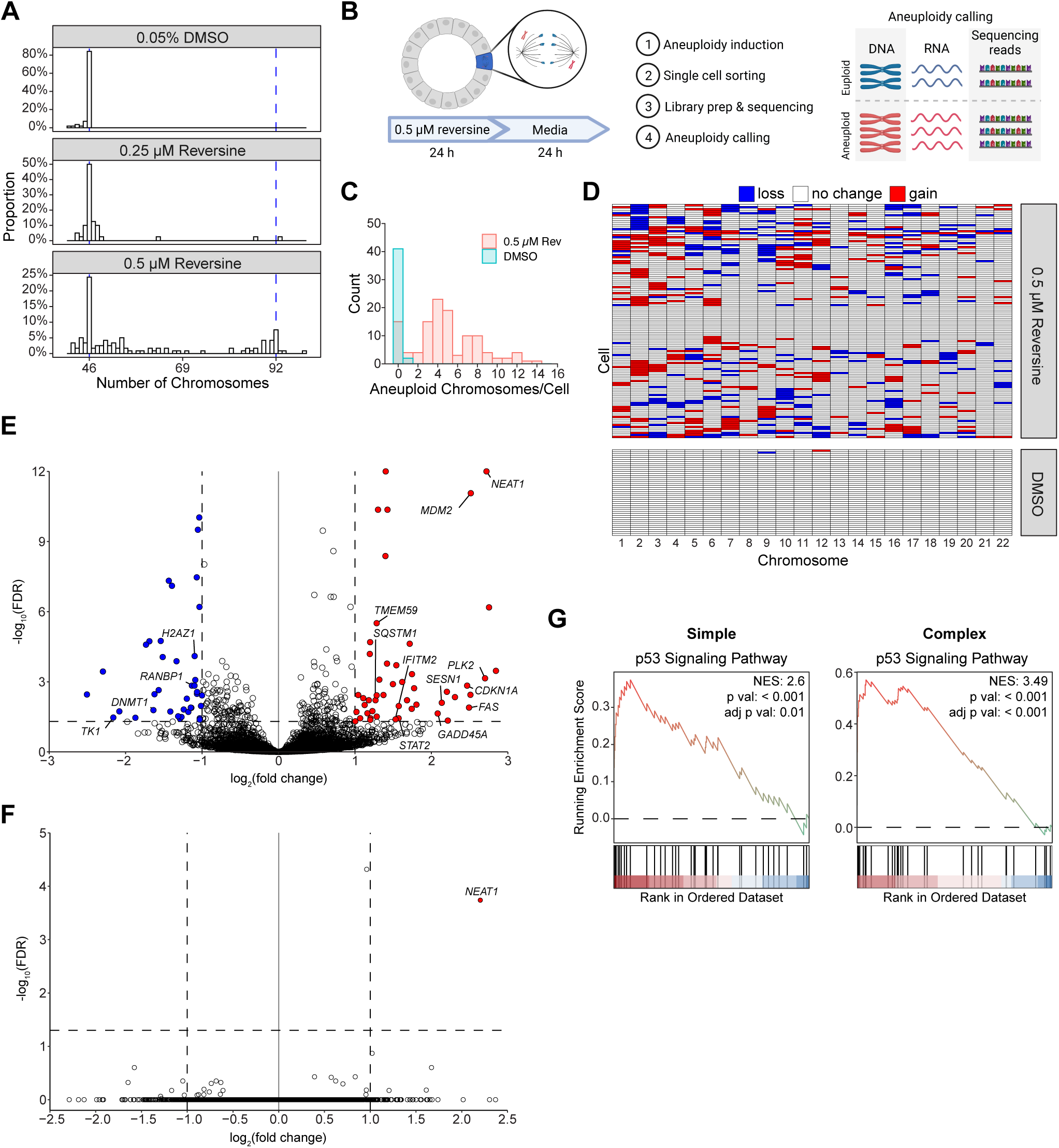
Common gene expression in complex aneuploid cells is associated with p53 activation. (A) Karyotype distributions assessed by metaphase spreads of colonoids treated with 0.05% DMSO, 0.25 µM reversine, or 0.5 µM reversine for 24 hours followed by 24 hours in drug-free proliferation media. n ≥ 3 independent experiments, n ≥ 30 metaphase spreads per experiment. (B) Schematic of single-cell RNA sequencing (scRNAseq) of aneuploid cells in colonoids. (C) Distribution of aneuploid chromosomes per cell from scRNAseq. Each chromosome gain or loss is defined as 1 aneuploid chromosome. (D) Karyotypes of 116 cells reversine treated and 45 vehicle control cells quantified from scRNAseq. Cells were grouped by hierarchical clustering. (E) Differentially expressed genes comparing complex aneuploid (defined as ≥ 4 aneuploid chromosomes) and euploid cells. Cutoffs for differential expression were log_2_(fold change) > 1 or < –1 (shown by vertical dashed lines) and FDR ≤ 0.5 (shown by horizontal dashed line). Genes with positive log_2_(fold change) are more highly expressed in complex aneuploid cells. (F) Differentially expressed genes comparing simple aneuploid (defined as ≤ 3 aneuploid chromosomes) and euploid cells. Features are the same as Panel E. (G) Gene set enrichment analysis plots demonstrating enrichment of genes in the p53 signaling pathway in simple or complex aneuploid cells. NES: normalized enrichment score.

### Common gene expression in complex aneuploid cells is associated with p53 activation

To investigate the common transcriptional consequences of aneuploidy, we performed high-depth, full-length single-cell RNAseq (scRNAseq) of cells from colonoids after treatment with 0.5 µM reversine (Figure 1B). 45 vehicle control treated and 116 0.5 µM reversine-treated cells were included in this analysis. Because the abundance of RNA transcripts across a chromosome scales proportionally with changes in DNA abundance,^15^ scRNAseq data has been used to infer chromosome copy number.^16–19^ Using this approach (see Methods), we detected aneuploid cells in colonoids treated with 0.5 µM reversine. The distribution of chromosome numbers was similar to that observed in bulk populations via chromosome counting and few aneuploid cells were detected within the vehicle-control treated colonoids (Figure 1C). The observed aneuploid karyotypes were heterogenous displaying both chromosome gains and losses across the genome (Figure 1D). Consistent with previous reports^20,21^, larger chromosomes were more likely to be mis-segregated (Figure S2A).

Differential expression analysis revealed increased expression of p53 responsive genes, including *NEAT1*, *MDM2*, *PLK2*, *CDKN1A*, *FAS*, and *GADD45A* in complex aneuploid cells relative to diploid cells (Figure 1E). Interestingly, the same comparison using the simple aneuploid population yielded only 1 of the above differentially expressed genes, *NEAT1* (Figure 1F). Gene set enrichment analysis also revealed strong enrichment of genes related to the p53 signaling pathway in complex aneuploid cells, with weaker enrichment observed in simple aneuploid cells (Figure 1G). Aneuploidy has previously been linked to immune modulation through cGAS-STING signaling and the senescence associated secretory phenotype (SASP).^22–24^ However, neither classical senescence markers, nor SASP associated genes, were among the enriched pathways in complex aneuploid cells (Figure S2B). Similarly, interferon stimulated genes associated with activation of cGAS-STING signaling were not upregulated in complex aneuploid cells (Figure 1E). Immune activation associated with JAK-STAT signaling was upregulated (Figure S2B). Oxidative phosphorylation related genes were enriched in simple aneuploid cells, suggesting possible metabolic remodeling (Figure S2C). Overall, these results demonstrate that the common transcriptional pattern across heterogenous complex aneuploid karyotypes is associated with the activation of p53 and that simple aneuploid cells are transcriptionally more similar to euploid cells.

### Karyotype complexity rather than DNA damage drives p53 activation in aneuploid cells

To confirm the p53 pathway was functional in our colonoids, we demonstrated that p21 abundance increased and colonoids failed to expand upon administration of Nutlin-3a (Figure S3A and S3B). Colonoids with complex aneuploidy induced by 0.5 µM reversine demonstrated increased abundance of p53 and p21, whereas neither protein was increased with simple aneuploidy (Figure 2A). Colonoids with complex aneuploidy also displayed impaired cell proliferation, marked by significantly reduced EdU incorporation,^25^ compared with euploid control or colonoids with simple aneuploidy (Figure 2B). DNA damage, which may result from chromosome mis-segregation or replication stress, has been proposed to lead to p53 activation in aneuploid cells.^9,10,23,26,27^ However, in colonoids expressing H2B-mNeon and mCherry-53BP1, DNA damage foci were infrequently observed following chromosome mis-segregation and did not increase following mitosis in the presence of 0.5 µM reversine (Figure 2C). To assess whether replication stress in aneuploid cells could cause DNA damage and lead to p53 activation, we treated colonoids with 0.25 or 0.5 µM reversine for 4 hours, followed by 16 hours in drug-free media to allow cells to progress through S phase. We did not observe any difference in pH2AX abundance between cells with or without increased p53 abundance, suggesting that DNA damage could not explain the observed difference in p53 (Figure S3C). We also blocked the DNA damage response both during and after aneuploidy induction using inhibitors of ATM and ATR, key kinases which link DNA damage signaling to p53 activation.^28^ We observed no change in p53 abundance, p21 abundance, or EdU incorporation (Figure 2D, S3D, 2E). These findings suggest DNA damage is not a major driver of p53 activation in complex aneuploid cells.

**Figure 2.**
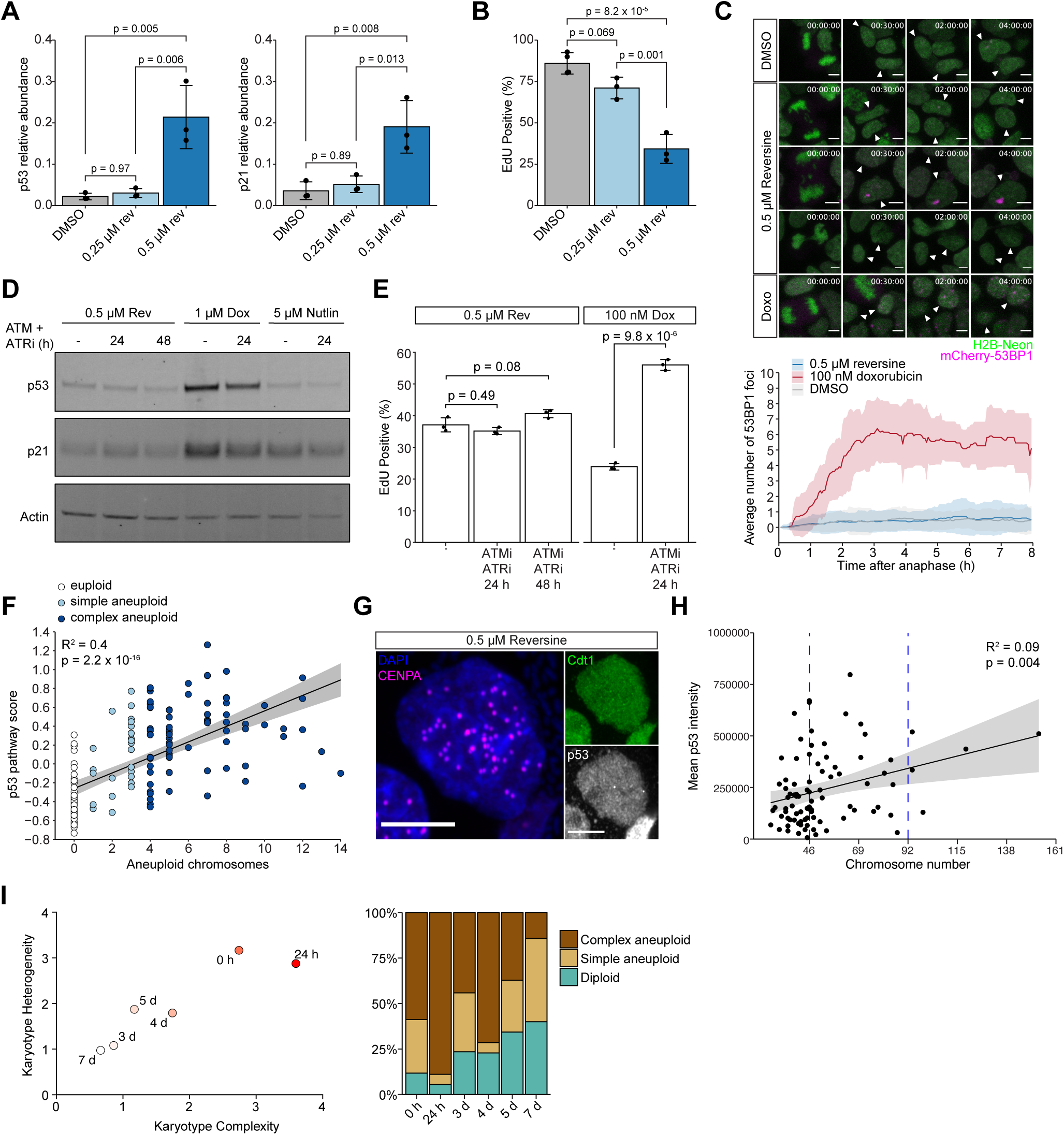
Karyotype complexity rather than DNA damage drives p53 activation in aneuploid cells. (A) p53 and p21 quantified by Western blot in colonoids treated for 24 hours followed by 24 hours in drug-free proliferation media. Relative abundance to GAPDH, loading control. n = 3 biological replicates. Error bars represent standard deviation. Abundances compared using ANOVA and post-hoc Tukey test. Tukey p values are shown. (B) Percentage of EdU positive cells quantified by flow cytometry. Conditions as in panel A. n = 3 biological replicates. Error bars represent standard deviation. Statistics as in panel A. (C) (top) Representative images from H2B-Neon-T2A-mCherry-53BP1 movies. (bottom) Quantification of 53BP1 foci following mitosis. Line represents the mean number of 53BP1 foci per cell and ribbon represents standard deviation. n = 3 biological replicates, n ≥ 50 cells for DMSO and reversine and n = 14 cells for doxorubicin. Scale bar = 5 µm. (D) p53 and p21 immunoblot with or without ATM (2 µM KU-60019) and ATR (2 µM VE-821) inhibition for 24 or 48 hours. n = 1 biological replicate. (E) EdU positive cells measured by flow cytometry. Conditions as in panel D. (F) Relationship between p53 pathway expression and the number of aneuploid chromosomes in cells treated with 0.5 µM reversine in scRNAseq data. Line shows linear regression. (G) Representative image of a fixed cell expressing Cdt1-eGFP-T2A-CENPA-Halo and immunostained for p53. The HaloTag ligand JF549 was added. Scale bar = 5 µm. (H) Relationship between chromosome number and mean p53 intensity in 0.5 µM reversine-treated cells from Cdt1-eGFP-T2A-CENPA-Halo + p53 immunofluorescence imaging. Line shows linear regression. n = 3 independent experiments, n = 87 total cells. (I) (left) Karyotype complexity and heterogeneity (see Methods) assessed by metaphase spreads of colonoids treated with 0.5 µM reversine for 24 hours followed by 0 hours, 24 hours, 3 days, 4 days, 5 days, or 7 days in drug-free proliferation media. (right) Percentage of diploid, aneuploid, or near tetraploid (88-96 chromosomes) metaphase spreads over time. n = 2 independent experiments

As p53 activation was much stronger in colonoids treated with the higher concentration of reversine, we examined whether p53 activation correlates with karyotype complexity on the single cell level. We observed that the p53 pathway expression score^29^ in individual cells from our scRNAseq dataset positively correlated with the number of aneuploid chromosomes. (Figure 2F). To directly visualize the relationship between chromosome number and p53 abundance, we generated colonoids expressing CENPA-Halo, a centromere marker,^30^ and eGFP-Cdt1, which allows visualization of G1 cells^31^ (Figure 2G). Using quantitative analysis to resolve clustered CENPA foci in G1 cells (see Methods), we identified cells with complex aneuploidy, although the error rate in counting prevented us from using this method to distinguish between simple aneuploid and euploid cells (Figure S3E). We observed a trend of increasing p53 immunofluorescence levels with increasing aneuploid chromosomes, although the correlation was just short of statistical significance (Figure 2H). Consistent with p53 and p21-induced cell cycle arrest in complex aneuploid cells, complex aneuploid karyotypes and overall karyotype complexity and heterogeneity decreased over 7 days following aneuploidy induction (Figure 2I, Figure S3F), but the overall frequency of aneuploidy remained greater than 50% with simple aneuploid karyotypes continuing to be prevalent.

### Direct tracking of proliferative cell fate following chromosome mis-segregation

To directly observe the fate of individual aneuploid cells, we generated colonoids expressing H2B-Dendra2, a photoconvertible fluorescent protein (FP), and converted one of the daughter nuclei from green to red fluorescence following chromosome mis-segregation (Figure 3A). Time-lapse imaging was performed to determine if the photoconverted cells progressed through the subsequent cell cycle (mitosis 1) (Figure 3B, Video S1). We observed that whereas mitotic errors in mitosis 0 reduced the frequency of observing a subsequent division (mitosis 1), daughter cells from divisions with chromatin bridges or monopolar spindles in mitosis 0 were less likely to divide again than those from divisions with lagging chromosomes (Figure 3C). We observed at most 3 lagging chromosomes in one division suggesting these daughter cells likely had simple aneuploid karyotypes. Consistently, a previous study demonstrated that divisions with lagging chromosomes frequently resulted in daughter cells with simple aneuploid karyotypes.^32^ Mitosis 0 errors did not delay cell cycle progression to mitosis 1 (Figure S4A).

**Figure 3.**
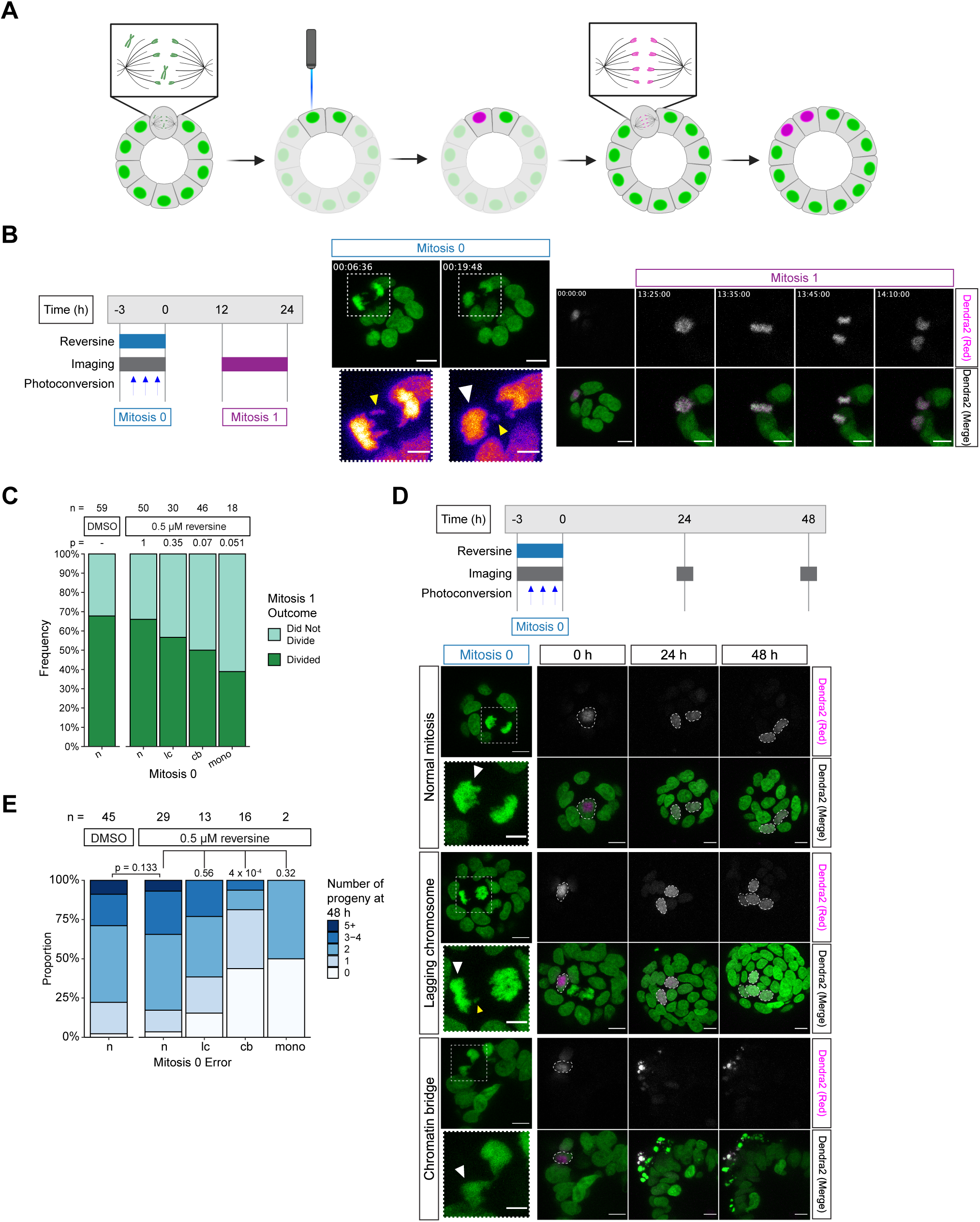
Direct tracking of proliferative cell fate following chromosome mis-segregation. (A) Schematic representation of H2B-Dendra2 photoconversion experiment. (B) (left) Schematic representation of experiment timeline. (right) Representative images of a photoconverted cell dividing following division with a lagging chromosome. White arrowhead denotes photoconverted cell. Yellow arrowheads denote a lagging chromosome. Scale bar = 10 µm, inset scale bar = 5 µm. (C) Division frequency of photoconverted cells in Mitosis 1. n ≥ 5 biological replicates. Total number of cells tracked is shown above each bar. p values calculated using Fisher’s exact test comparing condition of interest to DMSO. (D) (top) Schematic representation of experiment. (bottom) Representative images of photoconverted cells. Normal mitosis (top panel) was treated with 0.05% DMSO and lagging chromosome (middle panel) and chromatin bridge (lower panel) were treated with 0.5 µM reversine. White arrowheads denote the photoconverted cell. Yellow arrowhead denotes a lagging chromosome. Scale bar = 10 µm, inset scale bar = 5 µm. (E) Number of progenies 48 hours after Mitosis 0. n = 5 biological replicates. Total number of cells tracked is shown above each bar. p values were calculated using Fisher’s exact test.

We next quantified the number of progenies arising from a single photoconverted cell 48 hours (about 2 cell cycles) after chromosome mis-segregation (Figure 3D). Daughter cells arising from normal divisions or divisions with lagging chromosomes in mitosis 0 had a similar number of progenies, consistent with the lack of p53 activation or cell cycle arrest we observed in simple aneuploid cells (Figure 3E). Cells arising from chromatin bridges and monopolar spindles in mitosis 0 rarely divided further and the photoconverted FP was often fragmented and extruded from the epithelium suggesting cell death had occurred (Figures 3E, S4B). We found that chromosomes in daughter cells from divisions with chromatin bridges often coalesced with chromosomes from a neighboring nucleus during mitosis 1 indicating failed cytokinesis during mitosis 0 (Figure S4C). Consistent with previous findings,^33,34^ chromatin bridges can block cytokinesis leading to cleavage furrow regression (Figure S4D, Video S2).

To efficiently observe the effect of failed cell division on subsequent proliferation in colonoids, we directly induced tetraploid cells using ZM447439, MPI-0479605 with monastrol, or cytochalasin D as previously shown.^27,35^ Regardless of the drug(s) used, tetraploid cells were less likely to divide than their euploid or aneuploid counterparts (Figure S4E), confirming a robust mechanism exists for sensing and inhibiting proliferation of tetraploid cells in colonoids.^36^ When rare tetraploid cells did divide, they often underwent multipolar division (Figure S4F), which is known to generate complex aneuploid karyotypes.^37^ Thus, tetraploidy may contribute to the cell cycle arrest observed following treatment with 0.5 µM reversine both directly and through the generation of complex aneuploidy.

### Intestinal stem cells with simple aneuploidy exhibit impaired differentiation

As our results suggest that simple aneuploid cells can continue to divide, we further investigated the ability of these cells to maintain stem cell capacity and form colonoids from single cells. Simple aneuploid cells, induced by treatment with 0.25 μM reversine, retained expression of stem cell marker genes (Figure S5A). We found no difference in organoid-forming capacity between simple aneuploid and euploid cells (Figure 4A). However, colonoids generated from simple aneuploid cells grew more slowly (Figure 4B).

**Figure 4.**
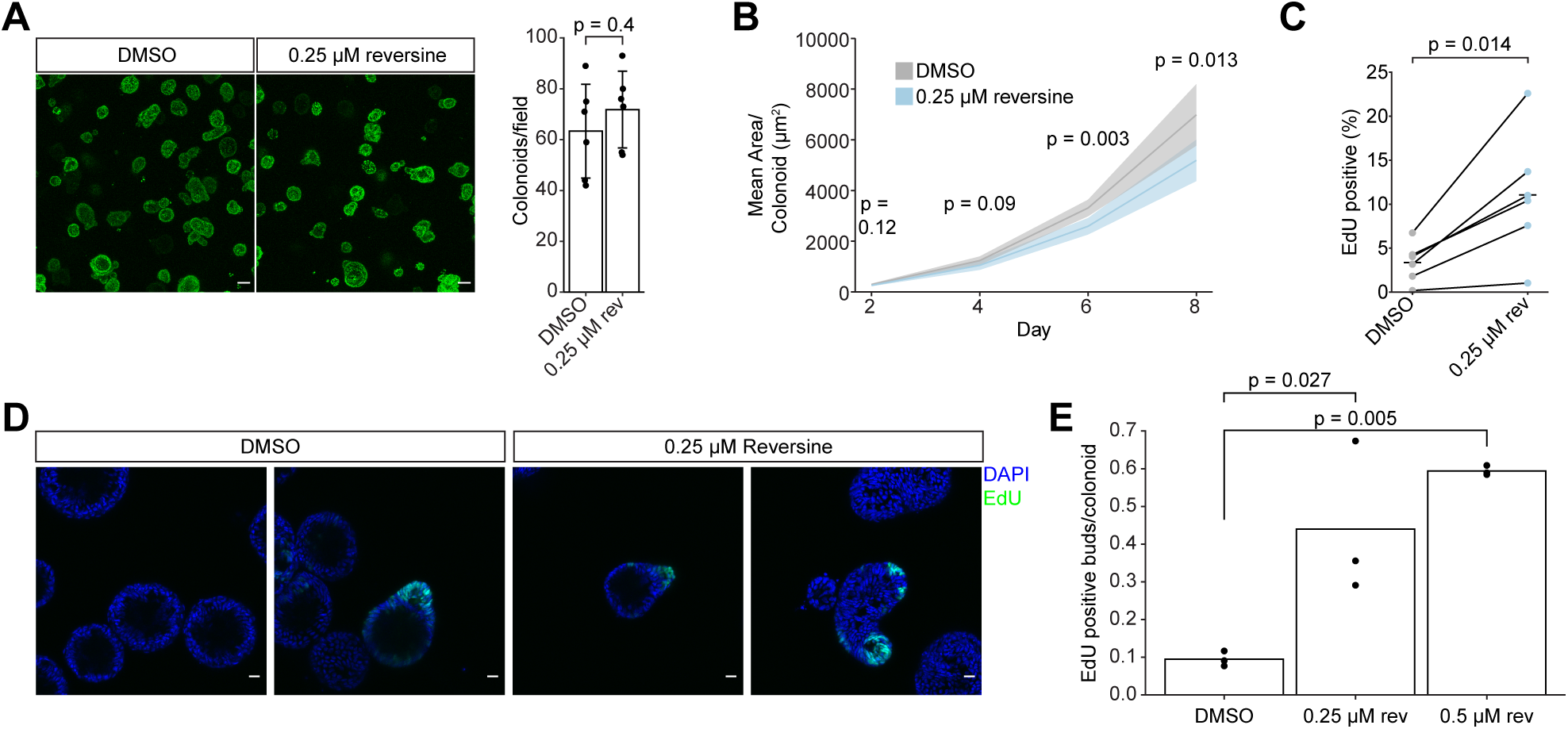
Intestinal stem cells with simple aneuploidy exhibit impaired differentiation. (A) (left) Representative images of H2B-Dendra2 colonoids formed from single cells 8 days after plating. (right) Quantification of the colonoid forming efficiency. n = 6 biological replicates. p value was calculated by Welch’s two sample t test. Scale bar = 100 µm. (B) Mean colonoid area over time of colonoids formed from single cells. Area was quantified from maximum intensity projections. n = 6 biological replicates. p values were calculated by Welch’s two sample t test. Ribbon shows standard deviation. (C) EdU positive cells in colonoids treated for 24 hours followed by 96 hours in drug-free differentiation media measured by flow cytometry. EdU was added to differentiation media for 16 hours. n = 5 biological replicates. p value was calculated using a paired t test. (D) Representative images of EdU localization in colonoid buds after 96 hours of differentiation. EdU was added to differentiation media for 16 hours. Scale bar = 20 µm. (E) Number of EdU positive buds per colonoid from images as in Panel D. n = 3 biological replicates. Compared using ANOVA and post-hoc Tukey test. Tukey p values are shown.

We then assessed the ability of aneuploid stem cells to differentiate, which can be tested in colonoids by removing stem cell growth factors and differentiation inhibitors from the culture media.^13^ We found that colonoids containing aneuploid cells were more likely to remain proliferative following 96 hours of differentiation compared to diploid controls (Figure 4C). As we did not observe higher expression of stem cell marker genes, as measured by RT-qPCR, in aneuploid colonoids compared with diploid controls grown in differentiation media (Figure S5B), impaired differentiation likely occurred in a minor fraction of the cells. Diploid colonoids became uniformly spherical following differentiation, whereas aneuploid colonoids contained crypt-like buds (Figure S5C). These buds frequently contained proliferative cells, suggesting that they were associated with stem cell niches (Figure 4D). Moreover, EdU-positive buds were more prevalent in aneuploid colonoids (Figure 4E), suggesting aneuploid colonoids retained more proliferative stem cells after differentiation. To rule out that slower proliferation of aneuploid colonoids delayed differentiation, we treated colonoids with low concentrations of Nutlin-3a or Palbociclib to slow down, but not arrest, growth (Figure S5D). These treatments did not increase colonoid budding (Figure S5E). Together, these findings suggest that simple aneuploid stem cells not only can continue to proliferate but also show impaired differentiation in the absence of stem cell niche factors.

## Discussion

Previous studies, using different experimental models, showed that the fate of aneuploid cells, especially p53 activation and cell cycle arrest, is varied and may be dependent on cellular and physiological contexts.^38–42^ Here, we investigated the effect of aneuploidy within the normal colon epithelium using patient-derived human colonoids. Using high-depth, full-length single cell RNA sequencing, biochemical assays, and live cell imaging approaches, our data suggest that the degree of aneuploidy plays a significant role in determining how cells respond. Cells with complex aneuploid karyotypes upregulated p53 and underwent cell cycle arrest or cell death. However, cells with simple aneuploid karyotypes did not mount a robust p53 response. Live cell tracking showed that cells with likely simple aneuploid karyotypes continued to divide. Notably, simple aneuploid cells are not benign as their presence led to increased propensity for stem cell niche factor-independent growth.

*TP53* mutations are strongly associated with aneuploidy across cancer genomes.^8,43,44^ However, more recent studies have questioned the role of p53 activation as a universal surveillance mechanism against aneuploidy, suggesting instead that p53 is activated in only a subset of aneuploid cells and often occurs indirectly as a result of DNA damage.^9–11,45^ By demonstrating an aneuploidy “dose-dependent” effect, our data supports the notion that chromosome imbalance plays a direct role in p53 activation. This is consistent with our previous work showing induction of mostly simple aneuploidy did not result in p53 activation in mouse colon or human mammary organoids.^12^ The underlying mechanism that leads to p53 activation remains unresolved, but two possible explanations may be explored in future research. First, p53 activation may be induced by a general aneuploidy signal or a common aneuploidy stress response. If the degree of such stress scales with the degree of chromosome imbalance, it could explain the association of p53 activation with complex aneuploidy. Second, there may not be a unified signal that leads to p53 activation, but instead the response may be driven by several different aneuploidy-associated stresses that converge on p53 activation. Many cellular stresses have been linked to aneuploidy and could contribute to p53 activation, including but not limited to DNA damage, replication stress, metabolic stress, immune activation, and hypo-osmotic-like stress.^10,23,24,26,46^ Our data, however, found that neither DNA damage nor replication stress are major contributors to p53 activation or cell cycle arrest in colonoids. It is possible that the DNA damage observed in cultured aneuploid cells in previous work^11,26^ is the result of environmental factors associated with 2-dimensional cell culture or the use of cell lines that have been passaged extensively *in vitro*.

The impact of aneuploidy on differentiation has been investigated in both embryonic and adult stem cells in a variety of organisms.^40,41,47–49^ Our data suggest that aneuploidy impairs differentiation of human colon stem cells, which have been demonstrated to be the cell of origin in colorectal cancer.^50^ Our finding is consistent with the observation that aneuploidy resulted in significant expansion of the proliferative zone in intestinal crypts in a mouse model of chromosomal instability.^42^ Although further studies are necessary to elucidate the underlying mechanism, this finding provides a plausible explanation for aneuploidy’s role in in the initiation of tumorigenesis, as niche-factor independent growth is an important early step in CRC evolution.^3^

## Acknowledgments

B.A.J., A.Z.L., and T.L. received support from the NIH Medical Scientist Training Program T32GM136577. This work was supported by Bloomberg Professorship funds to R.L. from Johns Hopkins University.

## Author Contributions

Conceptualization, B.A.J, A.N., and R.L.; Methodology, T.L., Y.W., and S.S.; Investigation, B.A.J, T.B, A.Z.L., Y.D., T.L., D.Z., Y.W., J.Z.; Resources, J.Z., T.C.L, S.S.; Writing – Original Draft, B.A.J and R.L.; Writing – Review & Editing, B.A.J, A.Z.L., J.Z., and R.L.; Visualization, B.A.J and J.Z.; Supervision, S.S, T.C.L, J.Z., and R.L.; Funding Acquisition, R.L.

## Data and code availability

scRNAseq data will be made publicly available via NCBI GEO database.

## Methods

Table. Key Resources Data.

Table. Key Resources Data
| REAGENT or RESOURCE | SOURCE | IDENTIFIER |
| --- | --- | --- |
| Antibodies |  |  |
| Anti-p53 [DO-1], mouse monoclonal | Abcam | Cat# ab1101, RRID:AB_297667 |
| Anti-p21 Waf1/Cip1 (12D1), rabbit monoclonal | Cell Signaling | Cat# 2947, RRID:AB_823586 |
| Anti-GAPDH (D16H11), rabbit monoclonal | Cell Signaling | Cat# 5174, RRID:AB_10622025 |
| Anti-β-actin (D6A8), rabbit monoclonal | Cell Signaling | Cat #8457, RRID:AB_10950489 |
| Anti-phospho-histone H2A.X (Ser-139), rabbit monoclonal | Cell Signaling | Cat #9718, RRID:AB_2118009 |
| Anti-rabbit IgG, HRP-linked Antibody | Cell Signaling | Cat# 7074, RRID:AB_2099233 |
| Anti-mouse IgG, HRP-linked Antibody | Cell Signaling | Cat# 7076, RRID:AB_330924 |
| Anti-mouse IgG (H+L), F(ab')<sub>2</sub> Fragment (Alexa Fluor® 488 Conjugate) | Cell Signaling | Cat# 4408, RRID:AB_10694704 |
| Anti-rabbit IgG (H+L), F(ab')<sub>2</sub> Fragment (Alexa Fluor 555 Conjugate) | Cell Signaling | Cat# 4413, RRID:AB_10694110 |
| Anti-mouse IgG (H+L), F(ab')<sub>2</sub> Fragment (Alexa Fluor 647 Conjugate) | Cell Signaling | Cat# 4410, RRID:AB_1904023 |
| Alexa Fluor 488 Polyclonal Antibody | ThermoFisher | Cat# A-11094, RRID:AB_221544 |
| Biological Samples |  |  |
| Human colon resection samples | This study |  |
| Chemicals, Peptides, and Recombinant Proteins |  |  |
| NMS-P715 | EMD/Millipore | Cat# 475949 |
| Nutlin-3a | Sigma Aldrich | Cat# N6287 |
| Nocodazole | Sigma Aldrich | Cat# M1404 |
| ZM447439 | Selleckchem | Cat# S1103 |
| Reversine | Cayman Chemical | Cat# 10004412 |
| Doxorubicin | Sigma Aldrich | Cat# AMBH324A4B72 |
| KU-60019 | Selleckchem | Cat# S1570 |
| VE-821 | Selleckchem | Cat# S8007 |
| GSK923295 | Selleckchem | Cat# S7090 |
| Cytochalasin D | Sigma | Cat# C8273 |
| Monastrol | Selleckchem | Cat# S8439 |
| MPI-0479605 | Selleckchem | Cat# S7488 |
| Palbociclib | Selleckchem | Cat# S4482 |
| KaryoMAX Colcemid Solution in HBSS | ThermoFisher | Cat# 15210040 |
| Janelia Fluor HaloTag Ligand 549 | Gift from Bin Wu lab |  |
| Critical Commercial Assays |  |  |
| NEBNext Single Cell Lysis Module | New England Biolabs | Cat# E5530S |
| NEBNext Single Cell/Low Input RNA Library Prep Kit | New England Biolabs | Cat# E6420L |
| NEBNext® Multiplex Oligos | New England Biolabs | Cat# E6440S |
| Click-iT™ EdU Alexa Fluor™ 488 | Invitrogen | Cat# C10420 |
| PureLink RNA Mini | Invitrogen | Cat# 12183018A |
| Superscript IV VILO Master Mix | Invitrogen | Cat# 11756050 |
| Fast SYBR Green Fast Mix | Applied Biosystems | Cat# 4385610 |
| Lenti-X concentrator | Takara Bio | Cat# 631232 |
| Recombinant DNA |  |  |
| pMD2.G | Laboratory of Didier Trono | RRID:Addgene_12259 |
| psPAX2 | Laboratory of Didier Trono | RRID:Addgene_12260 |
| pLV-H2B-Neon-T2A-mCherry-CAAX | This study |  |
| pLV-H2B-Neon-T2A-mCherry-53BP1 | This study |  |
| pLV-EGFP-Cdt1-T2A-CENPA-Halo | This study |  |
| pLV-H2B-Dendra2 | This study |  |
| Software and Algorithms |  |  |
| ImageJ | Schneider et al., 2012 | https://imagej.nih.gov/ij/ |
| R, version 3.6.1 | R Core Team, 2019 | https://www.R-project.org/ |
| RStudio, version 1.1.463 | RStudio Team, 2020 | http://www.rstudio.com/ |
| ggplot2, version 3.2.1 | Wickham, 2016 | https://ggplot2.tidyverse.org/ |

### Lead contact

Further information and requests for resources and reagents should be directed to and will be fulfilled by the lead contact, Rong Li.

### Patient-derived colon organoid culture

The human colon organoid line used in this study was generated from an ascending colon resection for CRC. All protocols were approved by the Johns Hopkins School of Medicine Institutional Review Board. The mucosa was separated from the underlying tissue and finely minced. To release crypts, EDTA was added to a final concentration of 31.25 mM and the minced tissue was placed on an orbital shaker at 160 rpm for 1 hour at 4°C. Crypts were isolated by differential centrifugation and embedded in Matrigel. Organoids were grown in 24 well plates (or 8-well chambered coverglass, Nunc LabTek II, catalog no. 155360 for live imaging and immunofluorescence experiments) in proliferation media containing 50% Wnt-3a conditioned media, 15% R-spondin-1 conditioned media, 10% Noggin conditioned media, EGF, B27 supplement, A-83-01, SB202190, and primocin. Differentiation media did not contain Wnt-3a, R-spondin-1, or SB202190. Organoids were passaged every 6-7 days by adding 1 ml 1x TrypLE Express directly to each well to dissociate organoids to single cells. Briefly, organoids were incubated at 37°C for 10-15 minutes with trituration every 5 minutes until a single cell suspension was generated. Cell number was counted using flow cytometry with propidium iodide staining to exclude dead cells. Single cells were embedded in Matrigel at a density of 600 cells/µl of Matrigel and 25 µl was added per well in 24 well plates or 15 µl per well in 8 well chambered coverglass. After Matrigel polymerized for 10 min at 37°C, 500 µl of media was added per well in 24 well plates or 400 µl per well in 8 well chambered coverglass. 10 µM Y-27632 was added to the media for the first 2 days after passaging. For modified organoids with puromycin resistance, 1 µg/ml puromycin was added to media.

### Single-cell RNA sequencing

For single-cell RNA sequencing, wild-type organoids were passaged as described above. 0.5 µM reversine or 0.05% DMSO was added for 24 hours on day 4 after passaging. Reversine and DMSO were then washed out and organoids were grown in proliferation media for 24 hours. Organoids were then dissociated to single cells by adding 1 ml 1x TrypLE Express directly to each well. Organoids were incubated at 37°C for 20 minutes with trituration every 5 minutes until a single cell suspension was generated. Cells were stained with ethidium homodimer to exclude dead cells. Single cells were then sorted directly into 96 well PCR plates containing 5 µl NEBNext cell lysis buffer with murine RNase inhibitor and flash frozen on dry ice. cDNA synthesis, amplification, and library generation were performed according to the NEBNext Single Cell/Low Input RNA Library Prep Kit manufacturer’s protocol. NEBNext multiplex oligos were used as index primers. the Biomek i7 Genomics Workstation. The libraries were sequenced on the NovaSeq 6000 system at PE150 with NovogeneAIT Genomics. Paired reads were then aligned with STAR and gene expression was quantified with HTSeq to generate a count matrix using Partek Flow software. Sequencing yielded a median of 7,742 genes detected and 4,212,297 total counts per cell. 29 cells with < 5000 genes detected or > 20% mitochondrial reads were excluded. 1 cell was excluded following dimensionality reduction. For inference of karyotype from scRNAseq data, we developed a custom pipeline to estimate chromosome-level changes in gene expression patterns which demonstrated reliable performance and higher accuracy compared to competing methods (in preparation). Briefly, normalized gene counts were summed across all genes on the same chromosome, which were then aggregated across all cells to estimate population median chromosome scores. For each cell in each chromosome, we calculated deviation from this median score and assigned gain (or loss) of chromosome as high (or low) chromosome score. We followed Griffiths et al. in gene expression normalization and chromosome score calculation^18^, followed by permutations of gene-chromosome relationships to estimate the significance of deviation for each (cell, chromosome) pair. Differential expression analysis was completed using DESeq2.^51^ Gene set enrichment analysis was performed using clusterProfiler.^52^ The p53 pathway score was calculated using the “AddModuleScore” function in the Seurat package.^53^ Genes used to calculate p53 pathway score were *CDKN1A*, *GADD45A*, *PLK2*, *MDM2*, *RPS27L*, *TRIAP1*, *FAS*.

### Lentivirus

Plasmids used: pMD2.G, psPAX2, pLV-H2B-Neon-ires-Puro, pLV-H2B-Neon-T2A-mCherry-CAAX-ires-Puro, pLV-H2B-Neon-T2A-mCherry-53BP1-ires-Puro, pLV-H2B-Dendra2-ires-Puro, pLV-eGFP-Cdt1-T2A-CENPA-Halo. HEK 293FT cells were co-transfected with the lentiviral transfer plasmid, packaging plasmid, and envelope plasmid. Media containing lentivirus was collected 24 and 48 hours after transfection. Lentivirus was concentrated using Lenti-X concentrator (Takara Bio, catalog no. 631232).

### Generation of modified organoid lines

Lentiviral transduction of organoids was performed as previously described ^54^. Organoids were dissociated to single cells and resuspended in 1850 µl organoid proliferation media, 50 µl concentrated lentivirus, 10 µM Y-27632, and 0.8 µg/ml polybrene. Cells were spinfected for 1 hour, 100 rpm, 28°C. Cells were incubated with virus at 37°C for 4-6 hours then rinsed to remove virus and embedded in Matrigel at a density of 1000 cells/µl Matrigel and grown in media containing 10 µM Y-27632 for the first 2 days after infection. Organoids were grown for 7 days and then passaged. 1 µg/ml puromycin was added immediately after the first passage.

### Drug treatments

The following drugs were used in this study: reversine, doxorubicin, nutlin-3a, NMS-P715, GSK923295, cytochalasin D, ZM44743, MPI-0479605, monastrol, nocodazole, KU-60019, VE-821, and palbociclib. Drug washout was performed by rinsing organoids three times with 1 ml PBS. Reversine was added 4 days after organoid passaging for all experiments using organoid proliferation media and 3-4 days after passaging for experiments involving organoid differentiation.

### Metaphase chromosome spreads

Cells were arrested in metaphase by adding 10 µg/ml Karyomax Colcemid for 4-6 hours. Organoids were then dissociated using TrypLE Express for 5-10 minutes at 37°C. Cell lines were collected using 0.05% trypsin dissociation. Cells were centrifuged at 300 x g, 5 minutes. The cell pellet was resuspended in 50 µl PBS then 5 ml 0.56% KCl solution was added for 13 minutes to promote cell swelling. 120 µl of 3:1 methanol:glacial acetic acid fixative solution was added and cell were pelleted by centrifuging at 300 x g, 5 minutes. Cells were resuspended in 7 ml 3:1 methanol:glacial acetic acid fixative solution and incubated overnight at 4°C. Cells were then dropped on glass slides and dried at 65°C for 20 minutes. Vectashield mounting media containing DAPI was added to each slide and a coverglass was placed and sealed. Spreads were imaged on a Nikon TiE-Eclipse epifluorescence microscope (60× oil immersion objective). At least 30 spreads were manually counted per condition. The absolute value of the difference of the chromosome number from the nearest euploid number (46, 92, 184) was calculated for each metaphase spread. This value was then divided by the nearest ploidy (2, 4, 8). The mean of these values for each condition was defined as the “karyotype complexity” and the standard deviation was defined as the “karyotype heterogeneity.”

### Live cell imaging

Organoids were plated on 8 well chambered coverglass (Nunc LabTek II, catalog no. 155360) for all live cell imaging experiments. Samples were imaged 4 days after seeding on a Zeiss 780 laser scanning confocal microscope with an environmental chamber maintained at 37°C with 5% CO_2_ using a 40x water immersion objective (NA 1.1). For imaging mitotic errors, organoids were imaged every 3-5 minutes with 15-20 2.0 µm z-slices and cell lines were imaged with 15-20 1.0 µm z-slices. Images were processed using a custom ImageJ macro modified from the ImageJ plugin “Temporal-Color Code.” Mitotic errors were quantified manually. For imaging 53BP1 foci, organoids were imaged every 3-5 minutes with 15-20 1.0 µm z-slices. The number of 53BP1 foci was counted manually.

### Photoconverting and tracking H2B-Dendra2

Organoids were seeded on 8 well chambered coverglass 4 days prior to imaging. 15 µl of Matrigel was added to each well at a density of 1200-1500 cells/µl of Matrigel. To bring all organoids into the same z-plane, the 15 µl of Matrigel was spread in a thin layer across the entire surface of the well. Organoids were grown in proliferation media supplemented with 10 µM Y-27632 and 1 µg/ml puromycin for the first two days. Reversine or vehicle-control was added 1 hour before the start of imaging. Total time of reversine treatment was limited to 4 hours. Cells were imaged and photoconverted using a Zeiss 780 laser scanning confocal microscope with an environmental chamber maintained at 37°C with 5% CO_2_ using a 40x water immersion objective (NA 1.1). Cells entering mitosis were manually identified and 12-16 cells were imaged per round of imaging with 2-3 rounds of imaging performed per experiment. Mitosis 0 was imaged every 3 minutes for a total of 24 minutes with 12 1.0 µM z-slices. The stage position of each mitosis was saved so the same cell could be imaged again later. After each round of Mitosis 0 imaging one daughter cells per observed mitosis was photoconverted from green to red. Photoconversion was performed using 10% 405 nm laser. An approximately 40 x 40 pixel region of interest was scanned 20 times with a pixel dwell time of 3 µsec. Each photoconversion took approximately 30 seconds. After all cells in that round were photoconverted, a 0-hour image on the entire organoid (green and red channels) was collected. After 2-3 rounds of Mitosis 0 imaging, each well was rinsed three times with PBS and 400 µl proliferation media was added to each well. For Mitosis 1 imaging, organoids were returned to the microscope 12 hours after the first round of Mitosis 0 imaging. Organoids were imaged every 7.5 minutes for 12 hours with 16 2.0 µm z-slices. For 24– and 48-hour imaging, organoids were returned to the microscope at 24 and 48 hours after the first round of Mitosis 0 imaging and each organoid was imaged then the slide was returned to the microscope. A custom ImageJ macro was used to process the images and movies to maximum intensity Z projections. Division during the Mitosis 1 imaging period and the number of photoconverted progeny in the 24– and 48-hour images were quantified manually.

### Western blot

Organoids were isolated by dissolving the Matrigel at 4°C in Cell Recovery Solution (Corning) with orbital shaking for 20 minutes. Cell lines were collected using 0.05% trypsin. Samples were pelleted by centrifugation (300 x g, 5 min) and protein was isolated immediately or the cells were flash frozen in liquid nitrogen. Cells were resuspended in 50-150 µl of lysis buffer containing 1X RIPA buffer, Roche Complete Mini, Roche PhoSTOP, 5% glycerol, 0.1% SDS. Cells were lysed by vortexing every 5 minutes for 25 minutes followed by sonication. Lysates were centrifuged at 16000 x g for 10 minutes at 4°C. Supernatants containing protein were collected and the protein concentration was measured using Bradford dye (BCA, Biorad). The lysate concentration was normalized and Bolt LDS Sample Buffer (Invitrogen, catalog no. B0007) and 40 mM DTT was added. Samples were separated in Bolt 4-12% Bis-Tris pre-cast polyacrylamide gels (Invitrogen). Proteins were transferred to PVDF membranes (Invitrogen, catalog no. IB24002) using the iBlot 2 Dry Blotting system (Invitrogen, catalog no. IB21001). Membranes were blocked with Intercept Blocking Buffer (LI-COR, catalog no. 927-60001) or 5% BSA in TBST (chemiluminescent blots). Primary antibodies were diluted in TBST and incubated with membranes overnight at 4°C with orbital shaking. Membranes were rinsed three times with TBST then secondary antibodies diluted in TBST were added for 1 hour at room temperature. Membranes were then rinsed one time with TBST followed by two times with TBS for fluorescent blots and three times with TBST for chemiluminescent blots. Membranes were then immediately imaged for fluorescent blot and imaged after addition of Clarity Western ECL substrate for chemiluminescent blots using a LI-COR Odyssey imager or Invitrogen iBright imager. Relative protein abundances were quantified using ImageJ. Quantification was normalized across biological replicates using sum normalization.

### EdU incorporation

Organoids were incubated with 10 µM EdU for 12 hours (Figure 2B) or 16 hours (all other EdU experiments). For flow cytometry, organoids were dissociated by adding 1 ml TrypLE Express to each well for 20-30 minutes at 37°C with trituration every 5 minutes until a single cell suspension was achieved. EdU labeling was performed using the manufacturer’s protocol for the Click-iT EdU Alexa Fluor 488 Flow Cytometry Assay Kit (Invitrogen, catalog no. C10420). EdU incorporation was measured by flow cytometry using the Attune flow analyzer. FCS files were analyzed using FlowJo. For imaging based EdU experiments, organoids were fixed with 4% PFA then permeabilized and blocked using the manufacturer’s protocol. Samples were imaged using a Zeiss 780 laser scanning confocal microscope with a 40x water immersion objective (NA 1.1). For quantification, budded organoids were identified based on a quantification of circularity. Then, the number of EdU positive buds was manually quantified in budded organoids.

### Immunofluorescence

Organoids were plated on 8 well chambered coverglass for immunofluorescence experiments. Media was removed and organoids were rinsed one time with PBS. Then, organoids were fixed in Matrigel with 4% PFA for 10 minutes at 37°C. The fixed organoids were rinsed three times with PBS for 10 minutes. Organoids were permeabilized with 0.5% Triton X-100 in PBS and blocked with 10% FBS and 0.1% Triton X-100 in PBS. Primary antibodies were diluted in 2% FBS and 0.1% Triton X-100 in PBS and added to samples overnight at 4°C. Organoids were then washed three times with PBS. Secondary antibodies diluted in 2% FBS and 0.1% Triton X-100 were added to samples for 2 hours at room temperature. Samples were rinsed one time with PBS then 1 µg/ml DAPI and/or 165 nM phalloidin was added for 10 minutes at room temperature and samples were rinsed one time with PBS. Samples were imaged using a Zeiss 780 laser scanning confocal microscope with a 40x water immersion objective (NA 1.1).

### Organoid formation assay

Organoids were dissociated by adding 1 ml TrypLE Express to each well for 20-30 minutes at 37°C with trituration every 5 minutes until a single cell suspension was achieved. Cell number was counted using flow cytometry with propidium iodide staining to exclude dead cells. Cells were embedded in Matrigel at a density of 600 cells/µl of Matrigel and 15 µl was plated per well on 8 well chambered coverglass. Organoids were grown in organoid proliferation media supplemented with 10 µM Y-27632 for the first 2 days. For fixed imaging, organoids were fixed with 4% PFA for 10 minutes at 37°C. Organoids were rinsed three times with PBS then permeabilized for 1 hour at room temperature with 0.5% Triton X-100 in PBS. Samples were rinsed one time with PBS then 1 µg/ml DAPI was added for 10 minutes at room temperature and samples were rinsed one time with PBS. For tracking of organoid growth in live cells, organoids were imaged every 2 days and positions were saved to image the same organoids each day. Samples were imaged using a Zeiss 780 laser scanning confocal microscope with a 10x objective for fixed cells and 20x objective for live cells.

### RT-qPCR

Organoids were collected by adding 500 µl TRIzol reagent (Invitrogen, catalog no. 15596026) directly to each well of organoids in Matrigel. This solution was flash frozen in liquid nitrogen and stored at –80°C until RNA extraction. RNA was extracted using the manufacturer’s protocol for the PureLink RNA mini kit (Invitrogen, 12183018A). RNA concentration was measured using a NanoDrop spectrophotometer. RNA was converted to cDNA using the Superscript IV VILO Master Mix (Invitrogen, catalog no. 11756050). RT-qPCR reactions were prepared using the Fast SYBR Green Fast Mix (Applied Biosystems, catalog no. 4385610). Quantification was performed using the ΔΔC_t_ method.

### General statistics

The number of biological replicates, the number of cells analyzed per biological replicate, and the statistical tests performed are indicated in the figure legends. A biological replicate was defined as a unique culture of organoids. Statistical tests were performed and figures were generated using R-Studio.

