## Supplemental Figures for "Differential effects of aneuploidy on growth and differentiation in human intestinal stem cells"

**Supplemental information titles and legends**

**Supplemental Video 1. Representative time-lapse H2B-Dendra2 tracking movie demonstrating division after chromosome mis-segregation, related to Figure 3.** Mitosis 0 time interval: 3 minutes 18 seconds, Mitosis 1 time interval: 5 minutes. Mitosis 1 imaging started 12 hours after mitosis 0. Non-photoconverted H2B-Dendra2 shown in green. Photoconverted H2B-Dendra2 shown in magenta. Mitosis 1 images were zoomed in to the photoconverted cell at the time of imaging.

**Supplemental Video 2. Representative time-lapse H2B-Dendra2 tracking movie demonstrating a binucleate daughter cell after mitosis with a chromatin bridge, related to Figure S4.**

Mitosis 0 time interval: 3 minutes, Mitosis 1 time interval: 5 minutes. Mitosis 1 imaging started 12 hours after mitosis 0. Non-photoconverted fluorescence shown in green. Non-photoconverted H2B-Dendra2 shown in green. Photoconverted H2B-Dendra2 shown in magenta. Mitosis 1 images were zoomed in to the photoconverted cell at the time of imaging.

**Supplemental Figure 1. Inducing aneuploidy in non-transformed human colonoids**

(A) Representative H&E stained tissue section from the colon resection sample used to generate the colonoid line used in this study.

(B) (left) Percentage of diploid (46 chromosomes), aneuploid, or near tetraploid (88-96 chromosomes) metaphase spreads following treatment with aneuploidy-inducing drugs. Colonoids were treated with drugs for 24 hours then metaphase spreads were performed after 24 hours of growth in proliferation media. At least 30 metaphase spreads were counted per condition. (right) Karyotype complexity and heterogeneity of each condition shown on left. See Methods for a discussion on the calculation of karyotype complexity and heterogeneity.

(C) Cumulative frequency plot of the nuclear envelope breakdown to anaphase onset time in human colon colonoids treated with 0.05% DMSO, 0.25 or 0.5 µM reversine. The time from NEBD-anaphase onset was calculated from live cell imaging of H2B-Neon or H2B-Neon-T2A-mCherry-CAAX colonoids. Combined data from ≥ 3 experiments.

(D) (left) Representative images of the mitotic errors. (right) Quantification of the frequency of mitotic errors observed in live cell images in human colonoids. Each dot represents the frequency of error in one experiment. Bars represent the mean of all experiments. $n \geq 2$ biological replicates and $n \geq 40$ divisions per condition. Fisher's exact test. *$p < 0.05$, **$p < 0.01$.

ZM = ZM447439, CEi = CENPE inhibitor (GSK923295), CytoD = cytochalasin D, NMS = NMS-P715, Rev = reversine, Noc = nocodazole

lc = lagging chromosome, cb = chromatin bridge, mono = monopolar

**Supplemental Figure 2. Single cell RNA sequencing chromosome distribution and pathway analysis**

(A) Bar plot showing the frequency at which each chromosome was gained or lost in all cells treated with 0.5 µM reversine quantified by single-cell RNA sequencing. Dashed line represents expected frequency of 25 aneuploid events per chromosome based on binomial($n$, $P$) distribution with $n = 116$ and $P = 0.21826$. *$p$ value $< 0.05$, **$p$ value $< 0.01$, *** $p$ value $< 0.001$.

(B) Gene set enrichment analysis of KEGG pathways utilizing all genes for complex aneuploid cells in scRNA-seq dataset.

(C) Gene set enrichment analysis of KEGG pathways utilizing all genes for simple aneuploid cells in scRNA-seq dataset.

**Supplementary Figure 3. Karyotype complexity determines p53 activation**

(A) Immunoblot of the human colonoid line used in this study treated with increasing concentrations of Nutlin-3a for 24 hours. GAPDH is a loading control.

(B) Images of cultures of the human colonoid line used in this study one week after passaging in the presence of increasing concentrations of Nutlin-3a.

(C) Quantification of standard deviation of pH2AX IF signal in p53 positive and p53 negative nuclei determined by IF in colonoids treated with DMSO, 0.25 µM reversine, 0.5 µM reversine, or 100 nM doxorubicin for 4 hours followed by 16 hours in drug-free media. n = 3 biological replicates (n = 2 for doxorubicin). Dots with color represent mean of each biological replicate. Unfilled dots represent individual nuclei.

(D) Bar graphs showing the abundance of p53 and p21 quantified by Western blot in colonoids treated with 0.5 µM reversine, 1.0 µM doxorubicin, or 5.0 µM Nutlin-3a for 24 hours with or without ATM and ATR inhibitors. Actin was used as a loading control and p53 and p21 abundance were quantified relative to GAPDH. n = 1 biological replicate.

(E) Karyotype distributions of chromosome numbers counted using CENPA-Halo foci. n = 3 biological replicates. n = 29 cell for DMSO, 22 cells for 0.25 µM reversine and 87 cells for 0.5 µM reversine. Blue dashed line at euploid and tetraploid chromosome numbers. Simple aneuploid cells shown between red dashed lines and complex aneuploid cells shown outside red dashed lines.

(F) Karyotype distributions assessed by metaphase spreads of human colonoids treated with 0.5 µM reversine for 24 hours followed by 0 hours, 24 hours, 3 days, 4 days, 5 days, or 7 days in drug-free proliferation media.

**Supplemental Figure 4. Chromatin bridges lead to binucleate cells that can undergo multipolar division**

(A) Cumulative frequency plot of the time from Mitosis 0 to Mitosis 1 in photoconverted H2B-Dendra2 cells.

(B) (left) Quantification of cell death in all photoconverted cells. The color indicates if cell death was first observed in the 24 hour or 48 hour image. (right) Quantification of cell death frequency for different mitotic errors in Mitosis 0. Quantified from the same images used for Figure 3D.

(C) Bar plot representing the frequency of all divisions in Mitosis 1 that were binucleate.

(D) Representative images of a chromatin bridge in Mitosis 0 that led to a binucleate, multipolar division in Mitosis 1. White arrowhead shows which daughter cell was photoconverted. Scale bar = 10 µm, inset scale bar = 5 µm.

(E) Division frequency of photoconverted tetraploid cells in Mitosis 1. $n \geq 2$ biological replicates. Number of cells tracked is shown above each bar. p values calculated using Fisher's exact test comparing condition of interest to DMSO in Figure 3C.

(F) Quantification of the frequency of bipolar and multipolar divisions in tetraploid cells that divided in Mitosis 1.

n = normal mitosis, lc = lagging chromosome, cb = chromatin bridge, mono = monopolar, ms = mitotic slippage, cf = cytokinesis failure, ZM = 1.0 µM ZM44743, CD = 0.75 µM cytochalasin D, M + M = 1.0 µM MPI-0479605 + 50 µM monastrol

**Supplemental Figure 5. Aneuploidy impairs intestinal stem cell differentiation**

(A) Expression of stem cell markers measured by qPCR over time following treatment with 0.05% DMSO or 0.25 µM reversine. DMSO and 0.25 µM reversine were washed out at time 0 hours and colonoids were grown in proliferation media for the remainder of the experiment. Points represent the mean of 3 biological replicates. Error bars represent standard deviation. p values were calculated using Welch's two sample t test.

(B) Expression of stem cell markers measured by qPCR over time following treatment with 0.05% DMSO or 0.25 µM reversine. DMSO and 0.25 µM reversine were washed out at time 0 hours and colonoids were grown in differentiation media for the remainder of the experiment. Points represent the mean of 3 biological replicates. Error bars represent standard deviation. p values were calculated using Welch's two sample t test.

(C) (left) Quantification of circularity in colonoids treated with 0.05% DMSO or 0.25 µM reversine then grown in drug-free proliferation or differentiation media with 96 hours. n = 3 biological replicates (right) Quantification of the frequency of colonoids that were budded after 96 hours in drug-free proliferation or differentiation media. colonoids were considered budded if their circularity was < 0.925. n = 3 biological replicates. p values were calculated using Fisher's exact test.

(D) Quantification of H2B-Dendra2 mean colonoid area over time. Drugs were added on Day -1 and drugs were washed out and differentiation media was added on Day 0. Line represents mean of 2 technical replicates.

(E) Quantification of circularity in colonoids treated with drugs shown for 24 hours then grown in differentiation media with 96 hours. n = 1 biological replicates. Each dot represents one colonoid. Line denotes mean circularity.

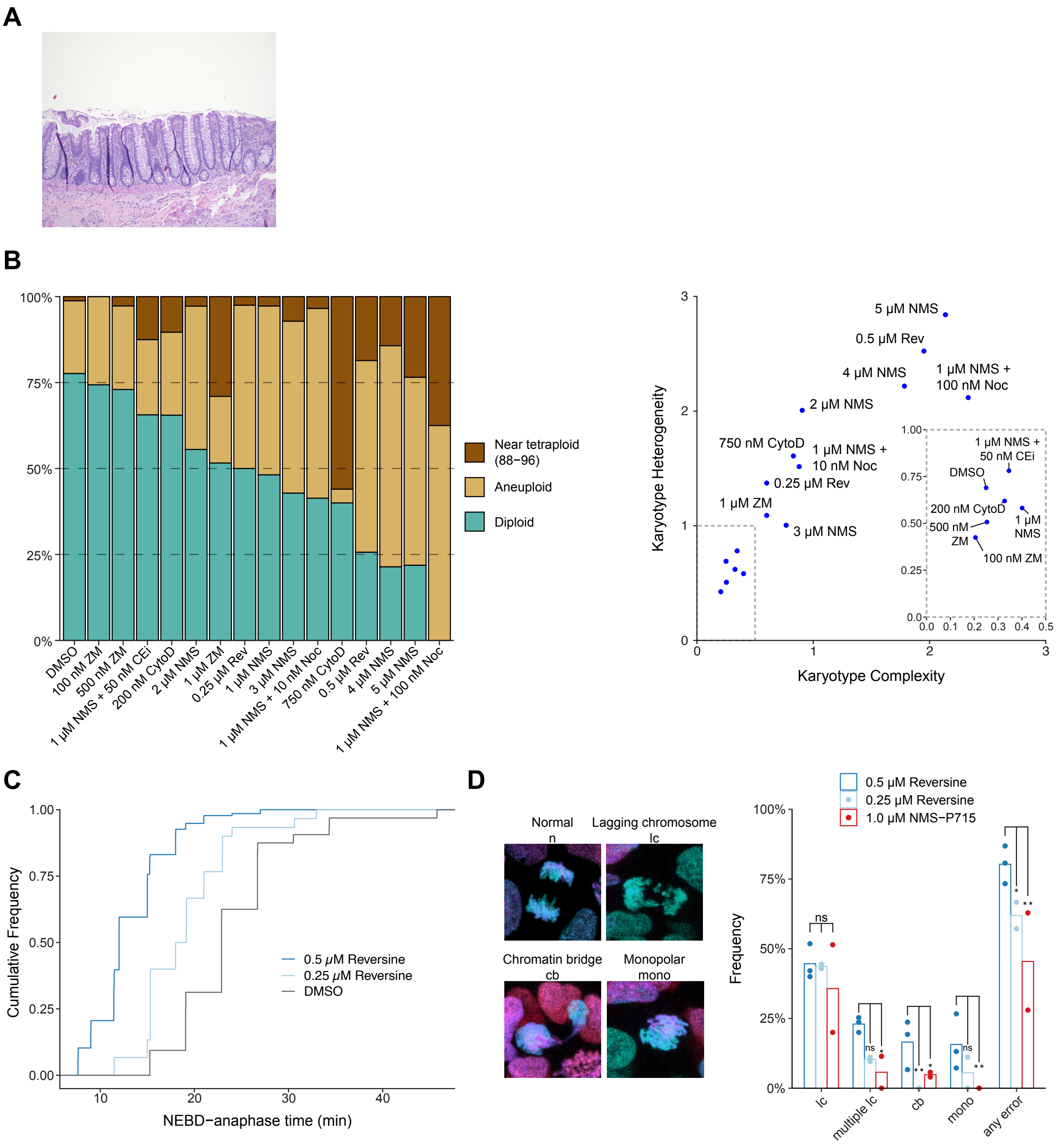


**Supplemental Figure 1. Inducing aneuploidy in non-transformed human colonoids**
(A) Representative H&E stained tissue section from the colon resection sample used to generate the colonoid line used in this study.
(B) (left) Percentage of diploid (46 chromosomes), aneuploid, or near tetraploid (88-96 chromosomes) metaphase spreads following treatment with aneuploidy-inducing drugs. Colonoids were treated with drugs for 24 hours then metaphase spreads were performed after 24 hours of growth in proliferation media. At least 30 metaphase spreads were counted per condition. (right) Karyotype complexity and heterogeneity of each condition shown on left. See Methods for a discussion on the calculation of karyotype complexity and heterogeneity.

(C) Cumulative frequency plot of the nuclear envelope breakdown to anaphase onset time in human colon colonoids treated with 0.05% DMSO, 0.25 or 0.5 μM reversine. The time from NEBD-anaphase onset was calculated from live cell imaging of H2B-Neon or H2B-Neon-T2A-mCherry-CAAX colonoids. Combined data from ≥ 3 experiments.

(D) (left) Representative images of the mitotic errors. (right) Quantification of the frequency of mitotic errors observed in live cell images in human colonoids. Each dot represents the frequency of error in one experiment. Bars represent the mean of all experiments. n ≥ 2 biological replicates and n ≥ 40 divisions per condition. Fisher's exact test. *$p < 0.05$, **$p < 0.01$.

ZM = ZM447439, CEi = CENPE inhibitor (GSK923295), CytoD = cytochalasin D, NMS = NMS-P715, Rev = reversine, Noc = nocodazole

lc = lagging chromosome, cb = chromatin bridge, mono = monopolar

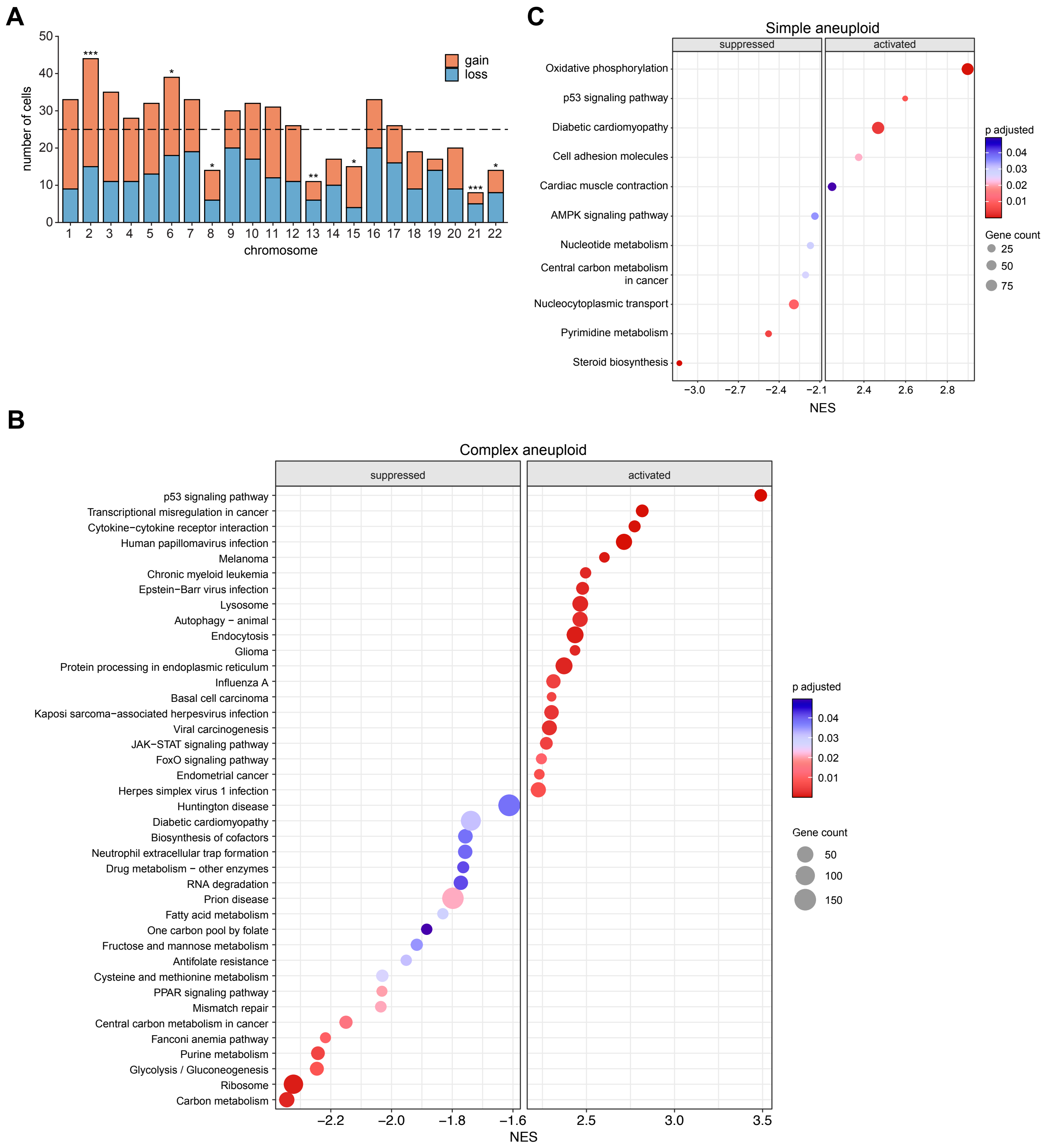


**Supplemental Figure 2. Single cell RNA sequencing chromosome distribution and pathway analysis**
(A) Bar plot showing the frequency at which each chromosome was gained or lost in all cells treated with 0.5 μM reversine quantified by single-cell RNA sequencing. Dashed line represents expected frequency of 25 aneuploid events per chromosome based on binomial(n, P) distribution with n = 116 and P = 0.21826. *p value < 0.05, **p value < 0.01, *** p value < 0.001.
(B) Gene set enrichment analysis of KEGG pathways utilizing all genes for complex aneuploid cells in scRNA-seq dataset. NES: normalized enrichment score
(C) Gene set enrichment analysis of KEGG pathways utilizing all genes for simple aneuploid cells in scRNA-seq dataset. NES: normalized enrichment score

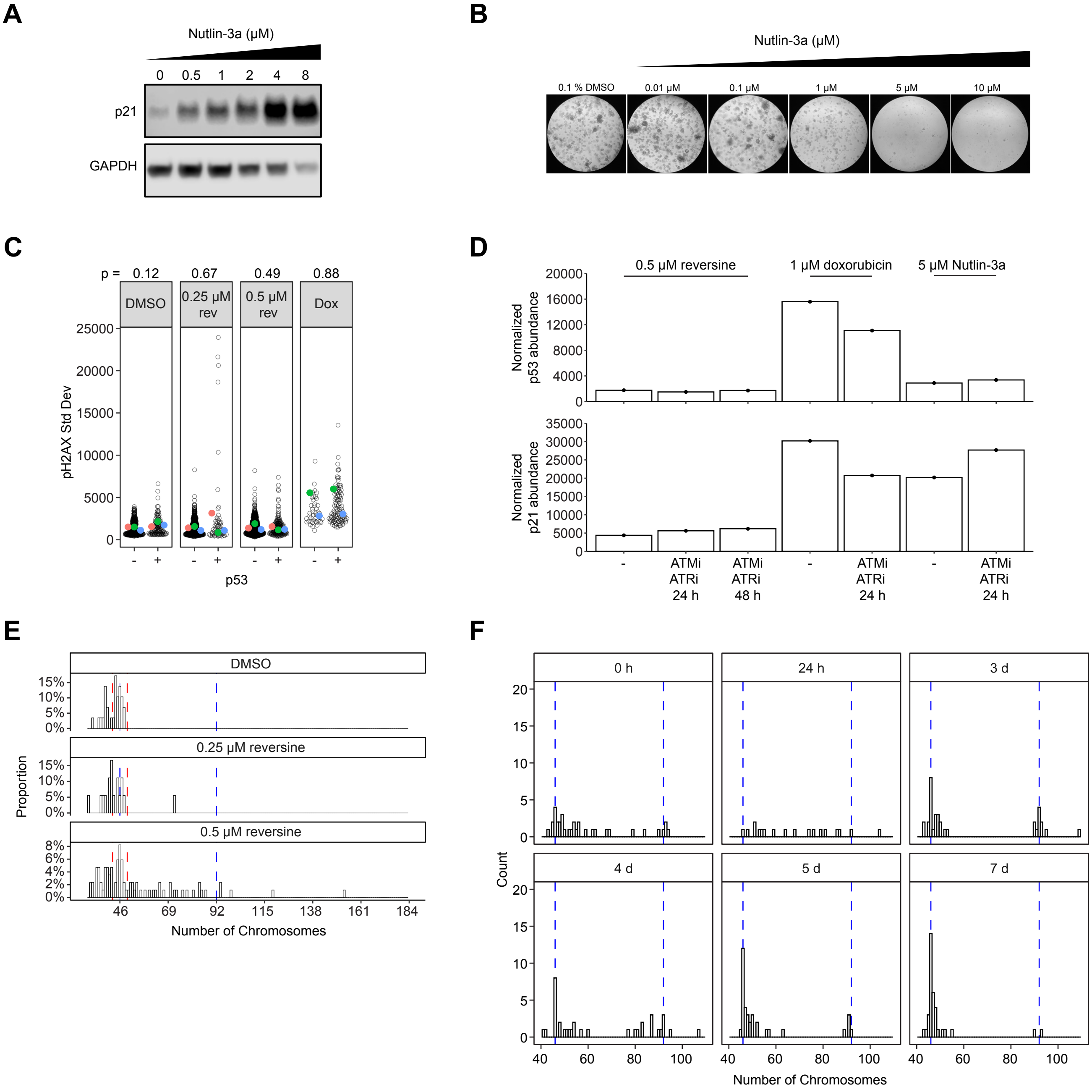


**Supplemental Figure 3. Karyotype complexity determines p53 activation**

(A) Immunoblot of the human colonoid line used in this study treated with increasing concentrations of Nutlin-3a for 24 hours. GAPDH is a loading control.

(B) Images of cultures of the human colonoid line used in this study one week after passaging in the presence of increasing concentrations of Nutlin-3a.

(C) Quantification of standard deviation of pH2AX IF signal in p53 positive and p53 negative nuclei determined by IF in colonoids treated with DMSO, 0.25 μM reversine, 0.5 μM reversine, or 100 nM doxorubicin for 4 hours followed by 16 hours in drug-free media. n = 3 biological replicates (n = 2 for doxorubicin). Dots with color represent mean of each biological replicate. Unfilled dots represent individual nuclei.

(D) Bar graphs showing the abundance of p53 and p21 quantified by Western blot in colonoids treated with 0.5 μM reversine, 1.0 μM doxorubicin, or 5.0 μM Nutlin-3a for 24 hours with or without ATM and ATR inhibitors. Actin was used as a loading control and p53 and p21 abundance were quantified relative to GAPDH. n = 1 biological replicate.

(E) Karyotype distributions of chromosome numbers counted using CENPA-Halo foci. n = 3 biological replicates. n = 29 cell for DMSO, 22 cells for 0.25 μM reversine and 87 cells for 0.5 μM reversine. Blue dashed line at euploid and tetraploid chromosome numbers. Simple aneuploid cells shown between red dashed lines and complex aneuploid cells shown outside red dashed lines.
(F) Karyotype distributions assessed by metaphase spreads of human colonoids treated with 0.5 μM reversine for 24 hours followed by 0 hours, 24 hours, 3 days, 4 days, 5 days, or 7 days in drug-free proliferation media.

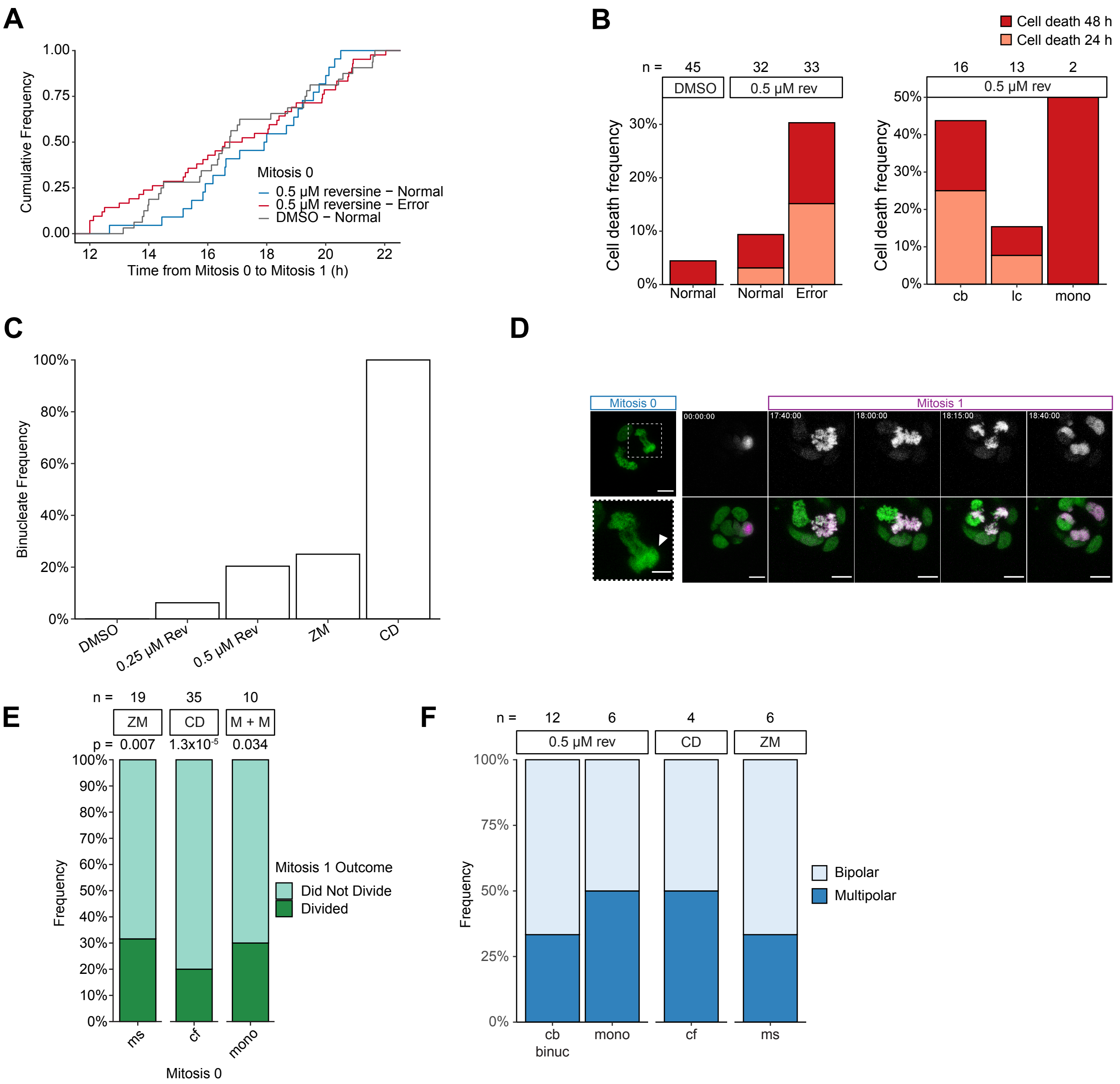


**Supplemental Figure 4. Chromatin bridges lead to binucleate cells that can undergo multipolar division**

A) Cumulative frequency plot of the time from Mitosis 0 to Mitosis 1 in photoconverted H2B-Dendra2 cells.

(B) (left) Quantification of cell death in all photoconverted cells. The color indicates if cell death was first observed in the 24 hour or 48 hour image. (right) Quantification of cell death frequency for different mitotic errors in Mitosis 0. Quantified from the same images used for Figure 3D.

(C) Bar plot representing the frequency of all divisions in Mitosis 1 that were binucleate.

(D) Representative images of a chromatin bridge in Mitosis 0 that led to a binucleate, multipolar division in Mitosis 1. White arrowhead shows which daughter cell was photoconverted. Scale bar = 10 μm, inset scale bar = 5 μm.

(E) Division frequency of photoconverted tetraploid cells in Mitosis 1. n ≥ 2 biological replicates. Number of cells tracked is shown above each bar. p values calculated using Fisher's exact test comparing condition of interest to DMSO in Figure 3C.

(F) Quantification of the frequency of bipolar and multipolar divisions in tetraploid cells that divided in Mitosis 1. n = normal mitosis, lc = lagging chromosome, cb = chromatin bridge, mono = monopolar, ms = mitotic slippage, cf = cytokinesis failure, ZM = 1.0 μM ZM44743, CD = 0.75 μM cytochalasin D, M + M = 1.0 μM MPI-0479605 + 50 μM monastrol

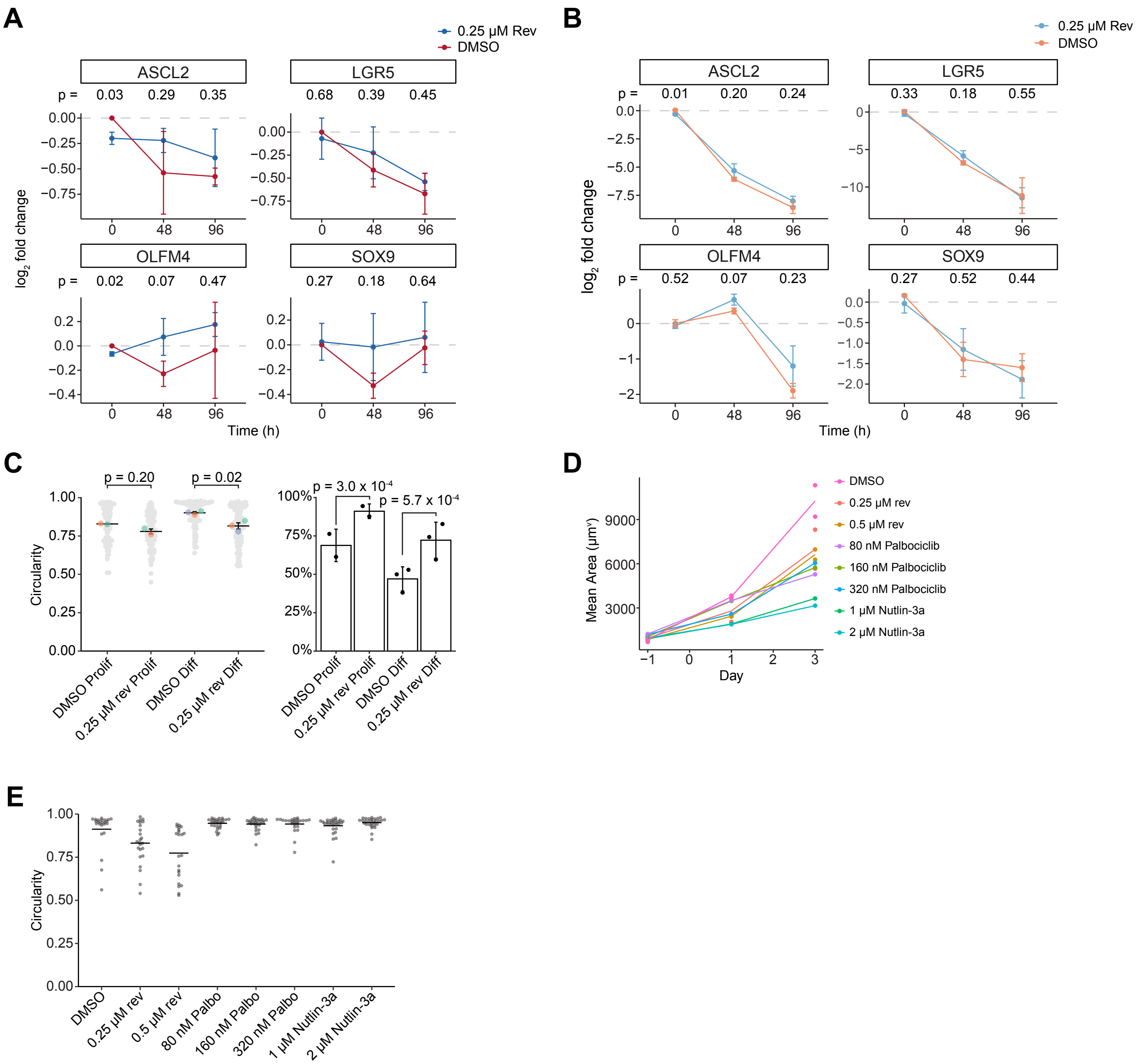


**Supplemental Figure 5. Aneuploidy impairs intestinal stem cell differentiation**

(A) Expression of stem cell markers measured by qPCR over time following treatment with 0.05% DMSO or 0.25 μM reversine. DMSO and 0.25 μM reversine were washed out at time 0 hours and colonoids were grown in proliferation media for the remainder of the experiment. Points represent the mean of 3 biological replicates. Error bars represent standard deviation. p values were calculated using Welch's two sample t test.

(B) Expression of stem cell markers measured by qPCR over time following treatment with 0.05% DMSO or 0.25 μM reversine. DMSO and 0.25 μM reversine were washed out at time 0 hours and colonoids were grown in differentiation media for the remainder of the experiment. Points represent the mean of 3 biological replicates. Error bars represent standard deviation. p values were calculated using Welch's two sample t test.

(C) (left) Quantification of circularity in colonoids treated with 0.05% DMSO or 0.25 μM reversine then grown in drug-free proliferation or differentiation media with 96 hours. n = 3 biological replicates (right) Quantification of the frequency of colonoids that were budded after 96 hours in drug-free proliferation or differentiation media. colonoids were considered budded if their circularity was < 0.925. n = 3 biological replicates. p values were calculated using Fisher's exact test.

(D) Quantification of H2B-Dendra2 mean colonoid area over time. Drugs were added on Day -1 and drugs were washed out and differentiation media was added on Day 0. Line represents mean of 2 technical replicates.

(E) Quantification of circularity in colonoids treated with drugs shown for 24 hours then grown in differentiation media with 96 hours. n = 1 biological replicates. Each dot represents one colonoid. Line denotes mean circularity.
